# Synthetic and genomic regulatory elements reveal aspects of cis regulatory grammar in Mouse Embryonic Stem Cells

**DOI:** 10.1101/398107

**Authors:** Dana M. King, Brett B. Maricque, Barak A. Cohen

## Abstract

In embryonic stem cells (ESCs), a core network of transcription factors establish and maintain the gene expression program necessary to grow indefinitely in cell culture and generate all three primary germ layers. To understand how interactions between four key pluripotency transcription factors (TFs), SOX2, POU5F1 (OCT4), KLF4, and ESRRB, contribute to cis-regulation in mouse ESCs, we assayed two massively parallel reporter assay (MPRA) libraries composed of different combinations of binding sites for these TFs. One library was an exhaustive set of synthetic cis-regulatory elements and the second was a set of genomic sequences with comparable configurations of binding sites. Comparisons between the libraries allowed us to determine the regulatory grammar requirements for these binding sites in constrained synthetic contexts versus genomic sequence contexts. We found that binding site quality is a common attribute for active elements in both the synthetic and genomic contexts. For synthetic regulatory elements, the level of expression is mostly determined by the number of binding sites but is tuned by a grammar that includes position effects. Surprisingly, this grammar appears to only play a small role in setting the output levels of genomic sequences. The relative activity of genomic sequences is best explained by the predicted affinity of binding sites, regardless of identity, and optimized spacing between sites. Our findings highlight the need for detailed examinations of complex sequence space when trying to understand cis-regulatory grammar in the genome.

## Introduction

Combinations of transcription factors (TFs) act at enhancers to specify cell states. Three models describe how TFs collaborate at enhancers, the billboard model, the enhanceosome model, and the TF collective model (Kulkarni and Arnosti 2003; Spitz and Furlong 2012). These models differ in the importance they ascribe to cis-regulatory grammar, defined as the extent to which the order, orientation, and affinity of transcription factor binding sites (TFBSs) impact the activity of enhancers. The enhanceosome model posits a strict grammar in which only precise arrangements of TFBSs activate target genes. The enhanceosome model is supported by structural studies of the IFN-β enhancer, where a specific order and spacing of TFBSs is required to activate expression (Yie, Senger, and Thanos 1999; Panne 2008). In contrast, the billboard model posits a more flexible grammar. In the billboard model enhancers can tolerate changes to the order, spacing, or orientations of TFBS with little change to target gene expression (Kulkarni and Arnosti 2003; Giorgetti et al. 2010) This model was proposed to explain binding site turnover in developmental enhancers and conservation in enhancer activity between species despite sequence divergence (Ludwig et al. 2000; Hare et al. 2008; Hare, Peterson, and Eisen 2008; Visel et al. 2009). In the TF collective model, specific TFs must be recruited to enhancers but can be recruited either by direct contact with DNA or indirectly through other TFs (Spitz and Furlong 2012; Junion et al. 2012; Uhl, Zandvakili, and Gebelein 2016). In the collective model no specific TFBS is required for activity even though individual TFs might be. These three cis-regulatory models differ in the importance that they give to TF-TF interactions and TF-DNA interactions in setting the activity of enhancers. The billboard model emphasizes the additive contribution of each TF bound to an enhancer, the enhanceosome model postulates geometrically constrained interactions between specific TFs, and the TF collective model highlights the joint action of TFs without requiring any particular TF-TF or TF-DNA interactions.

We and others have used mouse Embryonic Stem Cells (mESCs) as a system for studying cis-regulatory grammar and cooperative interactions between the pluripotency factors POU5F1 (OCT4), SOX2, ESRRB, and KLF4 (Fiore and Cohen 2016; Dunn et al. 2014; Williams, Cai, and Clore 2004). The pluripotency factors are a core set of TFs that maintain pluripotency in mESCs and are sufficient to induce pluripotency in terminally differentiated cells (Niwa 2014; Feng et al. 2009; Liu et al. 2008; Zhang et al. 2008; Takahashi and Yamanaka 2006). The pluripotency TFs activate self-renewal genes and repress genes that promote differentiation (Chambers and Tomlinson 2009). Based on known physical and genetic interactions, as well as genome-wide binding assays, multiple interacting TFs specify target gene expression in mESCs (Niwa 2014; Huang et al. 2009; Reményi, Schöler, and Wilmanns 2004; Williams, Cai, and Clore 2004; Reményi et al. 2003). However, it remains unclear how pluripotency TFs collaborate to drive specific patterns of gene expression in ESCs, and what role, if any, is played by TFBS grammar in determining target specificity in the genome (Chambers and Tomlinson 2009; X. Chen, Vega, and Ng 2008). Understanding how these factors combine to regulate their target genes is central to understanding the establishment and maintenance of the pluripotent state.

We previously addressed these questions by assaying a set of synthetic cis-regulatory elements that represent a small fraction of the possible arrangements of pluripotency TFBS. We identified some evidence for a grammar that is constrained by TFBS arrangement, including OCT4-SOX2 interactions. However our previous study lacked sufficient power to detect other interactions (Fiore and Cohen 2016). Here, we explore the role of grammar for pluripotency TFBSs by assaying an exhaustive set of synthetic cis-regulatory elements, composed of TFBS for SOX2, OCT4, KLF4 and ESRRB. The pattern of expression of synthetic regulatory elements is well predicted by a grammar that incorporates aspects of both the billboard and the enhanceosome models. However, this grammar seems to only play a small role in setting the activity of genomic regulatory elements with comparable configurations of binding sites. Genomic sequences appear to conform to a model where predicted TFBS affinity and favorable spacing between TFBSs contribute to activity levels, along with signals that recruit additional TFs, either directly through TF-DNA interactions or possibly indirectly through TF-TF interactions.

## Results

### Rationale and description of enhancer libraries

We designed two reporter gene libraries to explore the role of grammar in regulatory elements controlled by the pluripotency TFs. The first library, synthetic (SYN), is an exhaustive set of synthetic combinations of consensus TFBSs for OCT4 (O), SOX2 (S), KLF4 (K), and ESRRB (E). We did not include sites for NANOG in our libraries as its PWM has low information content and is not amenable to a synthetic binding site approach, in addition to being dispensable for reprogramming terminal cells to a pluripotent state (Jie Wang et al. 2013, 2012; Jauch et al. 2008; Pan and Thomson 2007; Takahashi and Yamanaka 2006). We also did not incorporate MYC binding sites in our libraries because MYC often acts independently of the core pluripotency TFs (C.-Y. Chen, Morris, and Mitchell 2012; Xi Chen et al. 2008; Liu et al. 2008). For each TF we used a binding site based on its Position Weight Matrix (PWM) in the JASPAR database (Sandelin et al. 2004; Fiore and Cohen 2016). We embedded each TFBS in a constant 20 bp sequence to ensure all the sites sit on the same side of the DNA helix (Fiore and Cohen 2016). We designed the SYN library to include all possible strings of two, three, and four TFBS building blocks (2-mers, 3-mers, and 4-mers, respectively), with each TFBS in either the forward or reverse direction, and each TFBS occuring no more than once per sequence, totaling 624 unique synthetic elements (Supplemental Table 1). The highly controlled nature of the SYN library provides maximum power to detect interactions mediated by the arrangement of binding sites.

The second library includes sequences from the mouse genome picked to match, as best as possible, members of the SYN library. Using the same PWMs used to design the SYN library, we scanned the mouse genome for combinations of the TFBSs for O, S, K, & E within 100 bp of regions bound by any of the four pluripotency TFs in E14 mESCs as measured by ChIP-seq (Fiore and Cohen 2016; Bailey et al. 2009; Xi Chen et al. 2008). We chose genomic sequences that contain one and only one binding site that scores above the PWM threshold for each factor to mimic the composition of the SYN library. We identified few clusters that included all four binding sites (< 70). We therefore selected 407 genomic sequences with three pluripotency TFBSs that could be compared to the exhaustive set of synthetic 3-mer elements. The resulting genomic wild-type library (gWT) is composed of the 407 unique genomic sequences with combinations of any three of the four TFBSs, with each site represented no more than once per sequence (Methods, Supplemental Table 3). Although these sequences differ from SYN elements in the individual site affinities, distances between TFBSs, as well as intervening sequence composition, our expectation was that the gWT sequences would directly test how well interactions learned from the SYN library apply to genomic sequences. To confirm that the activity of the gWT sequences depends on the presence of pluripotency TFBSs, we generated matched genomic mutant sequences (gMUT) in which all three of the identified pluripotency TFBSs were mutated by changing two positions in each TFBS from the highest information content base to the lowest information base according to the PWM (Supplemental Figure S7). The final gMUT sequences lack detectable TFBSs for O, S, K, or E when rescanned with the threshold used to select the gWT sequences. The combined gWT/gMUT library allows us to quantify the contributions of the pluripotency sites to regulatory activity, as well as sample configurations of pluripotency TFBS from the genome that may provide insight into grammar for these sequences.

### MPRA of reporter gene libraries

We assayed the cis-regulatory activity of the SYN and gWT/gMUT libraries in mESCs using a plasmid-based Massively Parallel Reporter Assay (MPRA) (Kheradpour et al. 2013; Mogno, Kwasnieski, and Cohen 2013; White et al. 2013; Patwardhan et al. 2012; Kwasnieski et al. 2012). Each unique library member described above is present eight times with a different unique sequence barcode (BC) in its 3’ UTR (Fiore and Cohen 2016; Mogno, Kwasnieski, and Cohen 2013; Kwasnieski et al. 2012). To determine the relative activation of each sequence compared to the minimal promoter included in each reporter construct, we included copies of plasmids with only the minimal promoter paired with over a hundred unique BCs in each library (See Methods) (White 2015; Fiore and Cohen 2016; Mogno, Kwasnieski, and Cohen 2013; White et al. 2013; Kwasnieski et al. 2012). Our measurements were highly reproducible between biological replicates, with R^2^ between 0.98-0.99 for replicates of the SYN library and 0.96-0.98 for the gWT/gMUT library (Figure S1A-B). After thresholding on DNA and RNA counts, we recovered reads for 100% (624/624) of our SYN elements and 99% (403/407) of paired gWT/gMUT sequences. The high concordance between replicates and simultaneous sequencing of the two libraries allowed us to make quantitative comparisons, both within and between libraries.

### Synthetic and genomic libraries support different grammar models

Synthetic regulatory elements have some characteristics of the billboard model. Most synthetic elements drive expression over basal activity regardless of the number, order, or orientation of sites within the element (Figure 1A). 77% of all SYN elements (6% of 2-mers, 66% of 3-mers, 92% of 4-mers) were statistically different from basal levels in all three replicates after correcting for multiple hypothesis testing (p < 0.05, Wilcoxon rank-sum test; Bonferroni correction, n = 637). In most cases, three or four consensus binding sites at fixed spacing are sufficient to increase expression above basal levels, which suggests strong independent contributions by binding sites in synthetic elements, as reported previously (Fiore and Cohen 2016). Elements with more binding sites generally drive higher expression than elements with fewer binding sites, which supports a billboard model of regulation, as the TFBSs can contribute to expression in an independent, additive manner. However, the wide range of expression observed for 4-mer elements (2.5-12.5, normalized expression) rules out a pure billboard model as differences between these elements must be due to the arrangement of the TFBSs, as site number and identity are fixed. The strong positive effect of adding sites supports a billboard model, while the diversity of expression of elements with the same number of sites reveals that grammar can quantitatively modulate activity.

**Figure 1.**
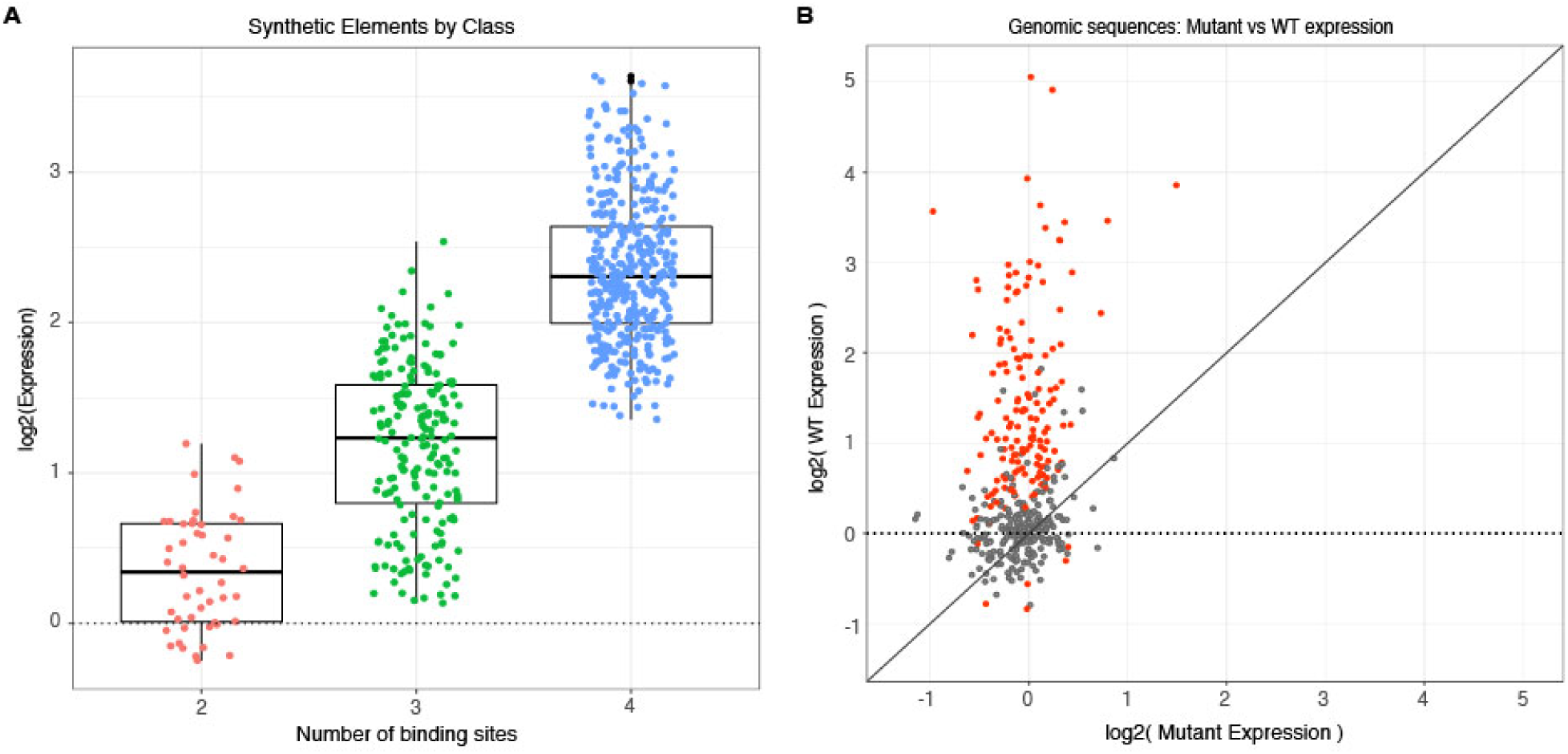
Activity of synthetic elements and genomic sequences. (**A**) The activity of synthetic elements with different numbers of binding sites. Expression is the average log of the ratio of cDNA barcode counts/DNA barcode counts for each synthetic element normalized to basal expression (dotted line). (**B**) The activity of genomic sequences is largely dependent on the presence of pluripotency binding sites. Normalized expression of wild type (gWT) sequences is plotted against expression of matched sequences with all three pluripotency TFBSs mutated (gMUT sequences). Red indicates sequences with significantly different expression between matched gWT and gMUT sequences (Wilcoxon-rank sum test, p < 0.05; Bonferroni correction, n = 403). Diagonal solid line is expectation if mutation of TFBSs had no impact on expression level. Expression of both gWT and gMUT sequences are normalized to basal controls, but basal expression is only plotted for gWT sequence on the y-axis (dotted line).

In contrast to the synthetic elements, most genomic sequences do not exhibit regulatory activity above basal levels. Only 28% (113/403) of wild type genomic sequences were statistically different from basal levels in all three replicates (p < 0.05, Wilcoxon rank-sum test; Bonferroni correction, n = 403). The low fraction of active gWT sequences demonstrates that three binding sites for the pluripotency TFs are often insufficient to increase expression above basal levels, a result that differs from the additive, billboard-like behavior observed for the SYN elements, but is consistent with observations from functional testing of genomic sequences bound by key TFs in other cell types (Fisher et al. 2012; Grossman et al. 2017; White et al. 2013). For genomic sequences that were statistically different from basal, 99% (112/113) have a significant difference between matched gWT and gMUT sequences (Figure 1B; p < 0.05, Wilcoxon rank-sum test; Bonferroni correction, n = 403), indicating that the activity of these sequences depends on one or more of the pluripotency TFBSs. Our observation that the presence of high-quality pluripotency TFBSs is generally insufficient to drive expression above basal levels does not support a strict billboard model for genomic sequences, at least when tested in a functional assay.

### Synthetic elements support a positional grammar

While the overall pattern of expression of SYN elements supports the billboard model, direct comparisons of different TFBS configurations also support a role for interactions between factors. Pairwise comparisons between a particular 3-mer and matched 4-mers that include one additional site at either the 5’ or 3’ end, reveal that the position of the extra site can strongly influence expression. For example, the O-K-E 3-mer and the matched O-K-E-<u>S</u> 4-mer drive indistinguishable expression, while the matched <u>S</u>-O-K-E 4-mer drives one of the highest expression levels in the SYN library (Figure 2A). Other examples are consistent with either strong position dependence or both position and orientation dependence (Figure S2A and S2B). Taken together these results show that when an additional TFBS is added to an existing synthetic element, the position and orientation of the new site can have large effects on activity.

**Figure 2.**
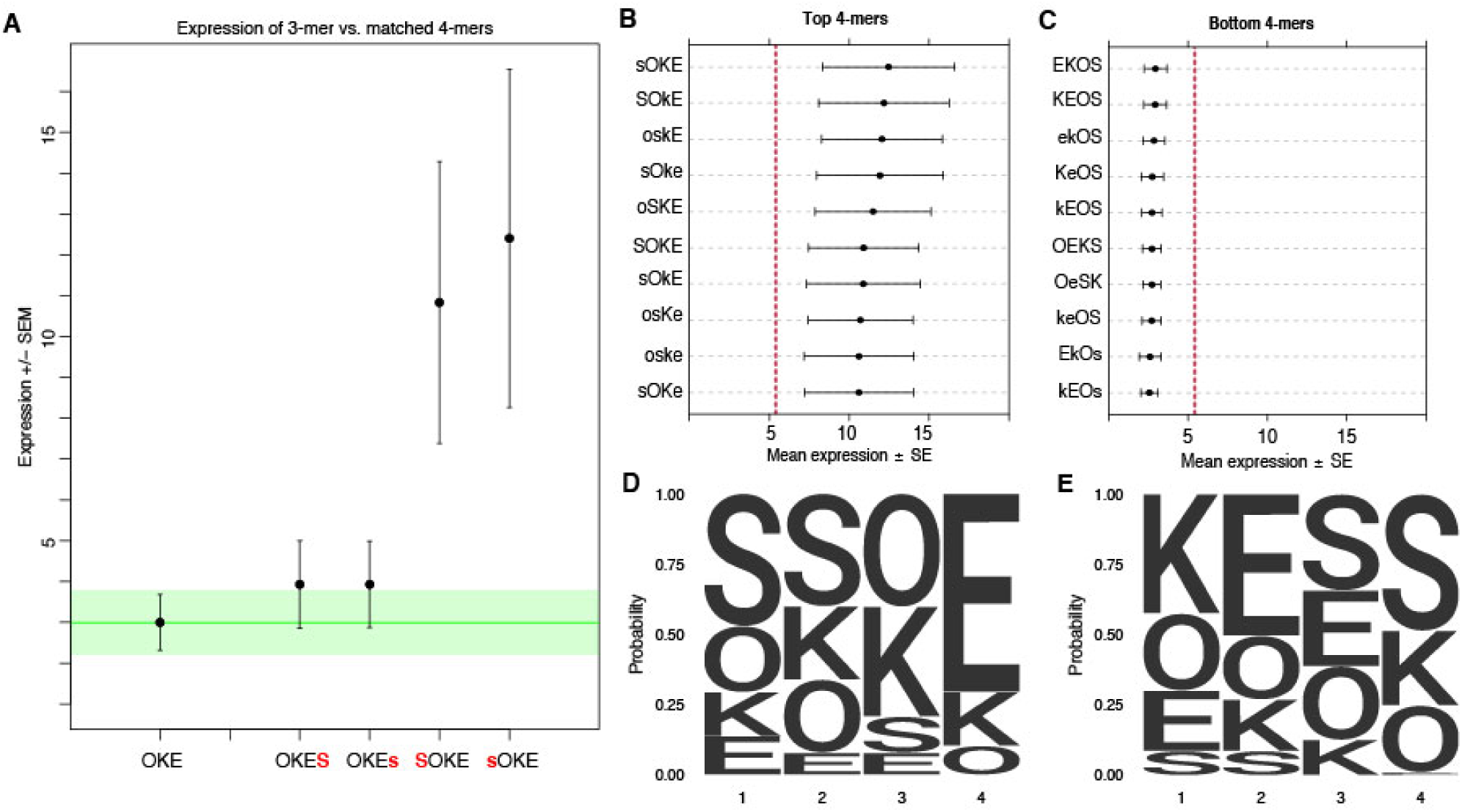
Non-additivity in synthetic elements. (**A**) Comparison of synthetic 3-mer elements with matched 4-mer elements containing one additional site in the first or fourth position. Mean expression of elements across barcodes (black dot) is plotted +/- SEM (black whiskers). Green line for comparison to expression of 3-mer; Green transparency highlights SEM of 3-mer shown. Activity of the ten highest (**B**) and ten lowest (**C**) expressing 4-mers. Red line represents average expression of all synthetic 4-mer elements. Capital letter represents binding site in forward orientation and lower-case letter represents binding site in reverse orientation. Mean expression of each element across barcodes (black dot) +/- SEM (black whiskers). Activity logos for the top 25% (n=96) (**D**) and bottom 25% (**E**) of 4-mer synthetic elements. Height of letter is proportional to frequency of site in indicated position. Positions organized from 5’ end (Position 1) to 3’ end (Position 4) of elements.

Synthetic elements appear to follow a grammar that includes position specific interactions between TFBSs. The ten highest expressing elements in the SYN library have a strong bias for S and O sites to be next to each other and in the first two positions (Figure 2B), while the ten lowest expressing 4-mers all have O and S in the last two positions (Figure 2C). The ten highest expressing 4-mers all have K followed by E in the last two positions, while the lowest expressing 4-mers tend to have K and E in the first two positions. We also found that the fourth position can have an especially large effect on expression. In the highest 25% of 4-mers (n=96) S is depleted (0/96) in the fourth position (Figure 2D), while in the lowest 25% E is virtually depleted (1/96) in the fourth position (Figure 2E). Conversely, in the fourth position, E is overrepresented in the top 25% (64/96) while S is overrepresented in the bottom 25% (48/96). These patterns also hold for comparisons of the strongest and weakest 3-mer and 2-mer elements (Supplemental Figures S2C-F). These patterns indicate a grammar that includes a positional bias to have adjacent SOX2 and OCT4 sites positioned upstream of KLF4 and ESRRB sites, which may favor interactions between these factors and the basal transcriptional machinery or TFs recruited by the minimal promoter. As specifying a site at a given position restricts possible sites in neighboring positions, these patterns could also represent favorable interactions between factors. The data suggest that synthetic elements have some properties consistent with the enhanceosome model, such as positional bias, with possible contributions by specific interactions between factors.

### Modeling supports role for TFBS positions in setting expression level for synthetic elements but not for genomic sequences

While the grammar of O, S, K, and E sites likely has an effect on the relative activities of the SYN elements, the grammar of these sites does not appear to contribute to the activity genomic sequences. We compared the SYN and gWT libraries for elements with configurations of OKE, OSE, OSK, and SKE TFBSs. Unlike SYN 3-mer elements, all four classes of gWT sequences span the range of expression observed for the entire library (0.56 to 33.2, normalized expression), with only OSK sequences having a higher average expression (Figure S3A). Thus, in genomic sequences, the same arrangement of sites embedded in different genomic contexts can either fail to drive detectable activity or drive expression higher than the highest SYN library member. To quantify the divergence in activities between genomic and synthetic elements directly, we matched gWT sequences with pluripotency TFBS dependent activity to SYN elements with the corresponding order of TFBSs. We observed no correlation in regulatory activity between matched site configurations, (R^2^ = 0.001; Figure S3B). These data indicate that other variables contribute to the cis-regulatory activity of gWT sequences, such as the spacing and affinities of the sites, or the presence of TFBSs for additional factors in flanking sequences that are held constant in the SYN library.

To identify additional sequence features that might be contributing to activity we used a variation of the Random Forest (RF) model, an unsupervised machine learning technique. RF models can be applied for either simple classification, assigning observations to group predictions, or classifying individual observations into semi-continuous bins to make quantitative, regression-case predictions. The accuracy of predictions are assessed over a large number of decision trees trained on random subsets of the data, which allows the contribution or “variable importance” of specific features to be measured. As RFs are prone to biases from early random splits in the decision trees for unbalanced data, we used iterative Random Forests (iRF) as a tool for feature selection as well as prediction (Basu et al. 2018).

We first trained a regression-case iRF model initialized with four features (Supplemental Table 4), representing only the presence or absence of each of the four pluripotency TFBSs to predict the expression of SYN elements. This “billboard” iRF model performed similarly to billboard-like thermodynamic based models trained on synthetic elements in mESCs (Fiore and Cohen 2016), with a R^2^ of 0.56 on a held-out test set for the final iRF iteration (2-fold cross-validation; Supplemental Figure S4). However, the billboard iRF model cannot account for the differences in activities between 4-mers, because all 4-mers have identical TFBS present (4-mers R^2^ = 0.00 (blue); Figure S4). To identify features that might distinguish between the activities of 4-mers, we trained an additional regression-case iRF model initialized with 20 features, representing both the presence and position of the four TFBS in each SYN element (Supplemental Table 4). The 20-term positional model performs well in predicting SYN expression, with an overall R^2^ of 0.87 for the last model iteration on a held-out test set (4-mers R^2^ = 0.52 (blue); Figure 3A). The positional iRF model highly weighs the presence/absence of the sites, as expected from the performance of the billboard iRF model, but also has contributions from the presence of ESRRB in the 4th position and SOX2 in the 1st and 2nd positions (Figure 3B). These results reinforce the conclusion that the activity of synthetic sequences depends both on the composition and positioning of TFBS.

**Figure 3.**
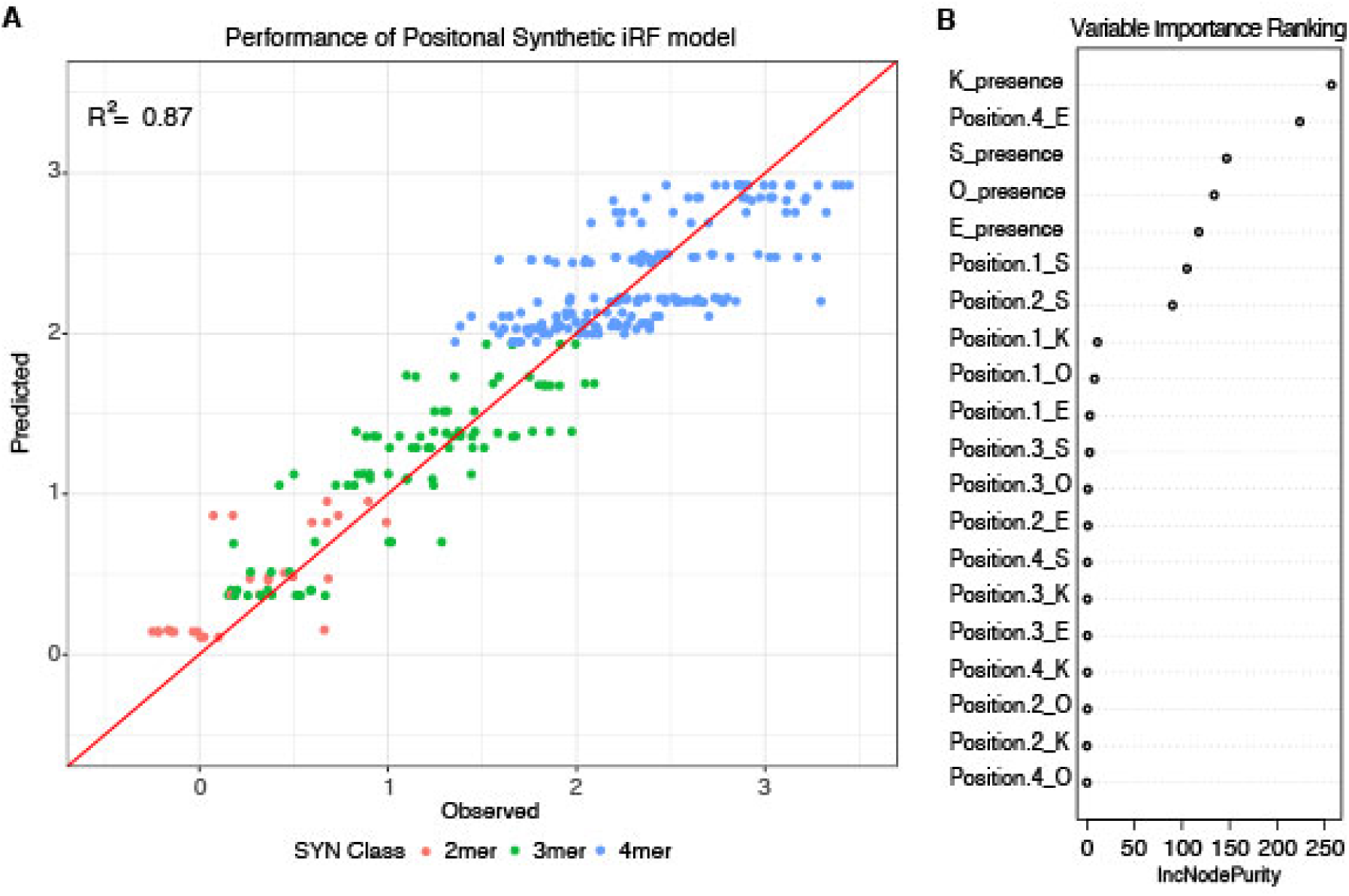
Positional grammar in synthetic elements. (**A**) Iterative random forest (iRF) regression model that includes features for presence and position of pluripotency TFBSs predicts relative expression of synthetic elements. Number of binding site per element is indicated in pink (2-mers), green (3-mers), and blue (4-mers). Observed and predicted expression are both plotted in log2 space. (**B**) Ranking of variables in synthetic iRF model. Variable importance is estimated by Increased Node Purity (IncNodePurity), the decrease in node impurities from splitting on that variable, averaged over all trees during training.

Models trained on the SYN library failed to predict or classify the expression of genomic sequences. While synthetic elements drove continuous expression across a range of activities, elements in the gWT library are predominantly inactive, and the small number of active gWT sequences drive expression across an order of magnitude of activity levels (dark green; Figure S3A). This presents a challenge for predicting gWT sequence activity with the features used to predict SYN elements. Retraining iRF regression models to predict gWT expression fails during the training step and has no correlation with the observed expression data (Billboard: R^2^ = 0.03; Billboard + Position: R^2^ = 0.001). However, training a classification model to distinguish between active and inactive gWT sequences (top 25%, n = 102; bottom 75%, n = 305) using either only billboard or billboard plus positional features also fails to perform better than chance (Billboard: AUROC = 0.52, AUPRC = 0.22; Positional: AUROC = 0.47, AUPRC = 0.25; Supplemental Table 5). When we directly compare the expression of gWT sequences to SYN elements with matching patterns of sites, we observe that not only do genomic sequences have a very different expression distribution from SYN elements, gWT sequences with the same pattern of TFBSs, as determined by motif matching, can drive drastically different expression levels (Supplemental Figure S3B). Other sequence features present in the genomic sequences and absent from the synthetic elements must therefore play a larger role in setting activity levels than the identity and position of the individual pluripotency TFBSs.

### Site affinity contributes to the activity of genomic sequences

We sought to identify features that differentiate active and inactive gWT sequences. Sequence-based support vector machines (kmer-SVMs) are powerful tools to predict the activity of putative regulatory elements independent of motif calling for specific factors that might be acting on the sequences (Fletez-Brant et al. 2013; Chaudhari and Cohen 2018). To identify sequence features that explain the differences between genomic elements, we trained a gapped kmer SVM (gkm-SVM) (Ghandi et al. 2016, 2014). The best performing gkm-SVM classified our positive and negative sets with AUROC of 0.75 and AUPRC of 0.77 (k = 8, gap = 2; Figure 4A). Although all sequences in the gWT library were selected to contain TFBSs for the four pluripotency factors, many of the discriminative 8-mers (29/50) have possible motif matches that include at least one pluripotency family member (Fletez-Brant et al. 2013; Bailey et al. 2009) (Supplemental File 6). This suggests that the differences between high and low activity genomic sites could be due to properties of either the primary pluripotency sites, defined here as TFBSs originally identified in the gWT library design, or due to secondary pluripotency sites, defined here as low PWM matches that scored below the scanning threshold, but could be present in the intervening sequences.

**Figure 4.**
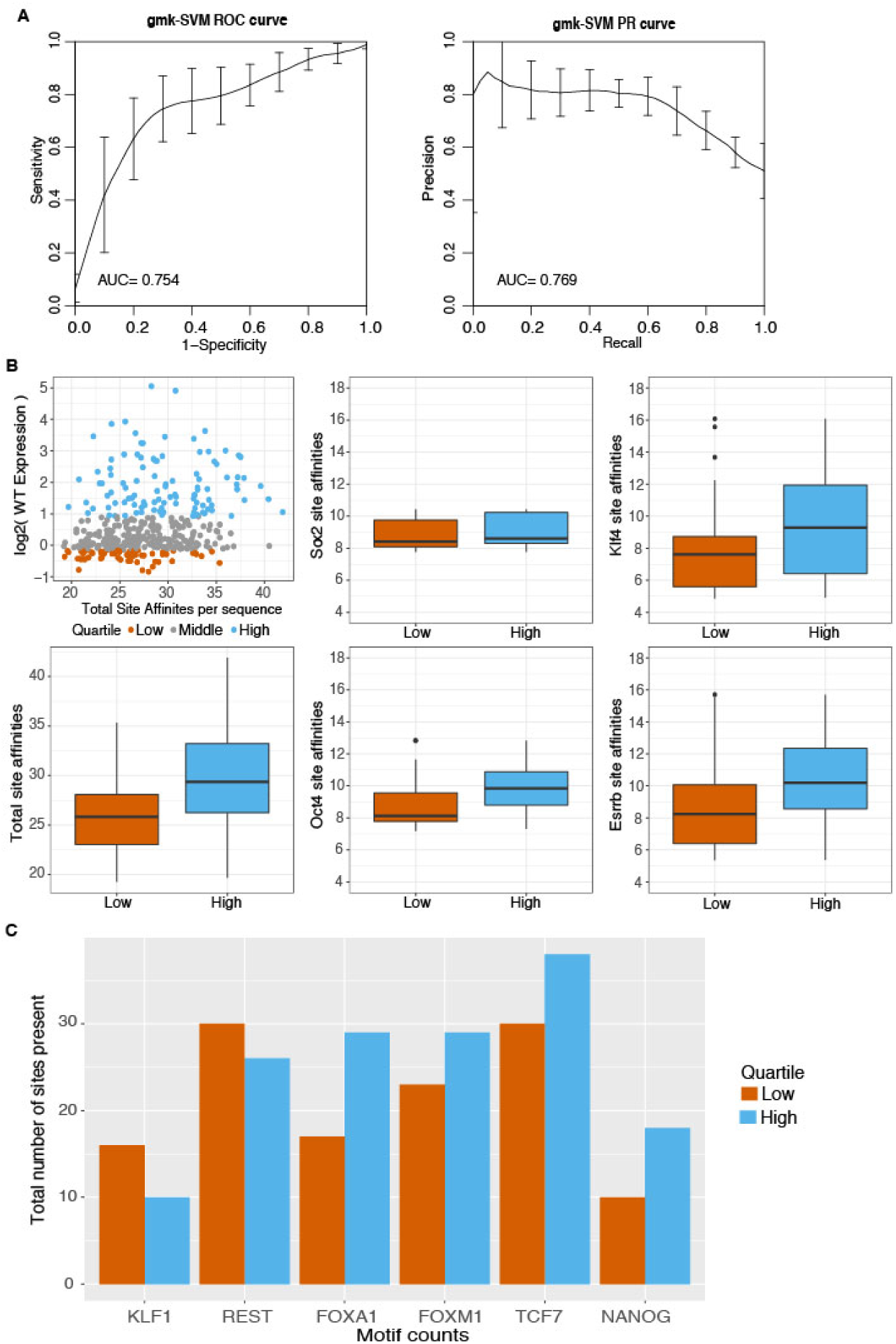
Sequence features separate active and inactive genomic sequences. (**A**) Performance of gkmer-SVM for genomic sequence supports contribution of sequence based features to activity. Word length of 8 bp with gap size of 2 bp was used for training with 3-fold cross validation. ROC curve (left panel) and PR curve (right panel) is plotted for the average across 3-fold cross-validation sets +/- standard deviation. (**B**) Primary (O,S,K,E) site affinities across gWT sequences, as output during motif scanning plotted for high genomic sequences (top 25% as ranked by expression, n = 101) and low genomic sequences (bottom 25% as ranked by expression, n =101). Total site affinities (bottom left panel) is calculated per sequences by summing predicted affinity of the three primary sites present in each sequence. (**C**) Total number of occurrences of TFBSs for additional TFs in high and low sequences (stratified as in **B**), as determined by motif scanning, excluding primary (O,S,K,E) sites.

Sequences with higher predicted affinity pluripotency TFBSs may drive higher expression. To determine if differences in the primary pluripotency sites are part of the signal identified by the SVM, we annotated gWT sequences with PWM-based scores for each TFBS present (Grant, Bailey, and Noble 2011). For SOX2, we found no difference in scores between high and low sequences (Figure 4B, panel 2; p = 0.07, Welch’s t-test). For OCT4, we found a modest difference between the average scores for high and low sequences and a broader but also a significant difference for KLF4 and ESRRB PWM scores (Figure 4B; p < 0.05, Welch’s t-test). The ‘OSKE_TotalAffinity’, the summed PWM scores for all four TFBS in each sequence, further separates high and low sequences (p < 0.05, Welch’s t-test). These patterns suggest that the quality of the primary sites contributes to the activity differences observed among gWT sequences.

We then asked if secondary sites for the pluripotency TFs might contribute to cis-regulatory activity by calculating predicted occupancy for both gWT sequences and gMUT sequences that lack the primary binding sites (See Methods). Predicted occupancy is a metric that includes contributions from any primary, well-scoring TFBSs plus contributions from weaker sites that might be missed with traditional motif scanning (White et al. 2016, 2013; Evans, Swanson, and Barolo 2012; Zhao and Stormo 2011; Segal et al. 2008). We found evidence for additional low predicted affinity sites for SOX2 and OCT4 in both high and low sequences, making it unlikely that low affinity sites strongly contribute to expression differences (Figure S5). Together, these results suggest that the affinities of the primary sites in genomic sequences, which are fixed in synthetic elements, contribute to the regulatory activity of genomic sequences more than the presence of additional sites with low predicted affinity.

### Contributions from sites for other transcription factors

A major difference between the synthetic and genomic elements is the presence of sites for TFs besides the pluripotency factors. While the synthetic elements were designed to keep the sequences between pluripotency sites constant, genomic sequences differ in both the length and composition of sequences between the pluripotency sites. The presence of binding sites for additional transcription factors may contribute to the activity of genomic sequences. To identify sites for other factors that could contribute to differences between high and low activity gWT sequences, we examined the top discriminative 8-mers from the gkm-SVM, looking at possible PWM matches for additional TFs (Supplemental File 6). We then used PWMs for these additional TFs to identify instances of sites for other factors in the genomic sequences (See Methods) (Grant, Bailey, and Noble 2011; Sandelin et al. 2004). We found significant enrichment for FOXA1 sites (Figure 4C; 1-sided Fisher’s exact test; 29 vs. 17 sites, p = 0.03, OR = 1.97). We also found that FOXA1 and NANOG had higher total PWM scores in the high activity sequences (Figure S6; 1-sided Welch’s t-test; FOXA1: p = 0.04; NANOG: p = 0.03). While FOXA1 is likely not present in mESCs, other family members (FOXA2, FOXD1, FOXP1) are expressed in ESCs and have been shown to contribute to the pluripotent regulatory network (Pan and Thomson 2007; Mulas et al. 2018; Gabut et al. 2011), and therefore could be acting on the gWT sequences through these binding sites.

Genomic sequences with higher occupancy by TFs in the genome, as measured by ChIP-seq, have higher average expression in our assay. We annotated the gWT intervals with publicly available ChIP-seq data for additional TFs and with ATAC-seq data from E14 mESCs to determine differences in accessibility (Supplemental Table 4). We found that accessibility was generally high for both high and low activity gWT sequences, and therefore not enriched. High activity sequences were significantly enriched for overlap with NANOG peaks (Supplemental Figure S7; 1-sided Fisher’s exact test, NANOG: p = 0.03, OR = 2.17). However, for the 328 genomic sequences with a NANOG ChIP-seq signal, only 16% had an underlying TFBS as determined by motif scanning. Therefore, NANOG might be recruited by other pluripotency TFs to these sequences independent of high quality TFBS for this factor. If we compare expression levels to the number of overlapping ChIP-seq peaks, including O,S,K,E and these additional TFs, we see that gWT sequences with higher occupancy in the genome have higher average expression in our assay (Figure 5), which has been previously observed in HepG2 cells (Ulirsch et al. 2016).

**Figure 5.**
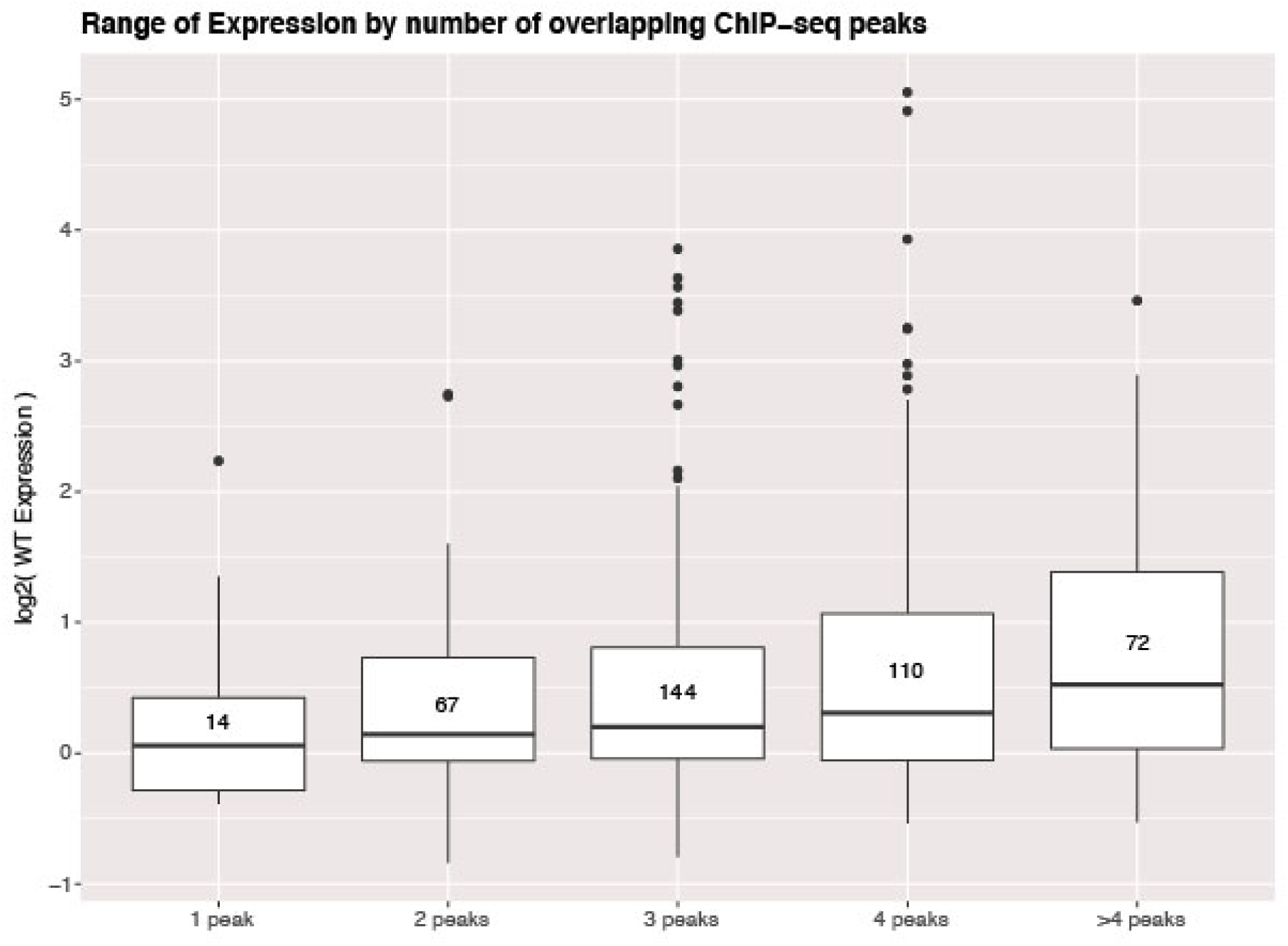
Activity of genomic sequences scales with increased occupancy in the genome. Expression of elements binned by number of intersected ChIP-seq peak signals for different factors. Number of sequences in each bin indicated in center of boxplot. All gWT sequences overlapped at least one ChIP-seq peak as per library design.

To understand the relative contributions of the features that had some predictive power on their own, we trained iRF models with subsets of these features and compared the performance of these models on a held-out test set (Supplemental Table 4). Including parameters only related to spacing between TFBSs, which is held constant in SYN elements but variable in gWT sequences, resulted in a model with an AUROC of 0.52 (Figure 6A) and AUPRC of 0.31 (Figure 6B) (model ‘Spacing’). A model that includes parameters describing the attributes of the primary pluripotency sites, such as the scores for the sites, yields an AUROC of 0.64 and AUPRC of 0.34 (model ‘PrimarySites’). We then limited our model to only parameters that relate occupancy of these regions by TFs in the genome, as measured by ChIP-seq, which resulted in a model with an AUROC of 0.59 and AUPRC of 0.31 (model ‘ChIP’). We trained a final iRF model initialized with 58 features that capture key differences between gWT sequences and SYN elements. These features include predicted affinity and spacing between the pluripotency TFBSs, predicted occupancy for the pluripotency TFs, binding sites for additional TFs, plus chromatin accessibility (ATAC-seq) and ChIP-seq peaks for both TFs and histone marks, as well as summary features such as the total primary site affinities (as in Figure 4B) for each sequence (See Supplemental Table 6 for full list of features). This gWT iRF model performed fairly well on a held out test set (AUROC = 0.67, AUPRC = 0.46; model ‘All’). The features that best separate active sequences were related to attributes of the pluripotency sites with the top feature being the summed pluripotency factor predicted affinity per sequence (‘OSKE_TotalAffinity’, Figure 6C). Taken together our data suggests that genomic sequences are able to drive higher expression when they contain stronger predicted affinity binding sites for pluripotency TFs with optimal spacing and are embedded in sequences that can mediate the recruitment of other TFs or cofactors.

**Figure 6.**
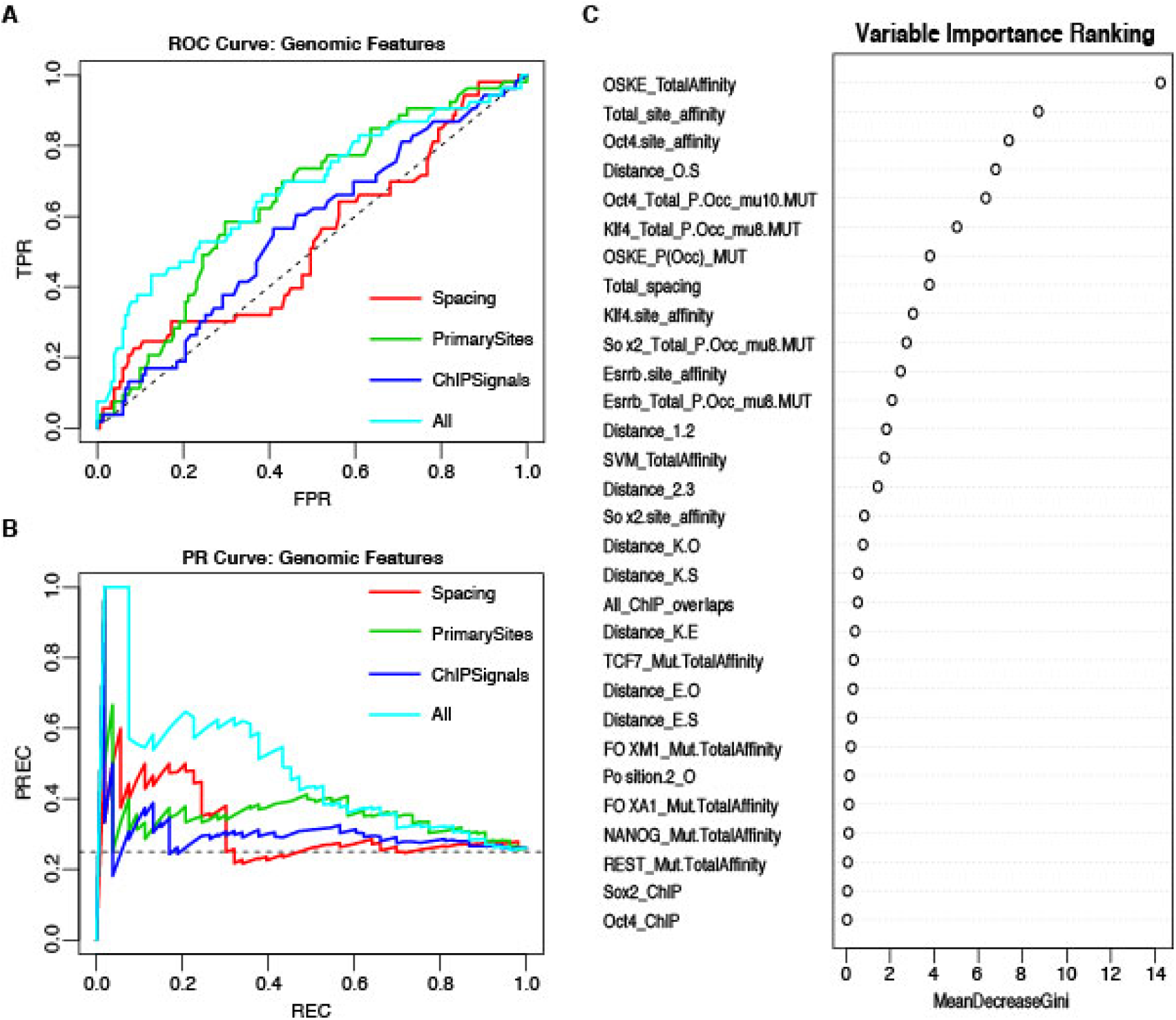
Performance of iRF classification models that include features specific to genomic sequences. (**A**) ROC Curve and (**B**) Precision-Recall (PR) Curve comparing genomic iRF models. Color indicates set of features used to train model. (**C**) Variable importance as evaluated for the feature by the average reduction in the Gini index, which is based on node impurity during training (Xi Chen and Ishwaran 2012).

## Discussion

In this study we sought to understand how pluripotency factors collaborate to drive specific levels of expression by testing both an exhaustive set of synthetic arrangements of TFBSs for OCT4, SOX2, KLF4, AND ESRRB and comparable genomic sequences. Our experimental design allowed for direct comparisons between the regulatory potential and regulatory grammar of synthetic elements and genomic sequences. Using a massively parallel reporter assay (MPRA), we found that the regulatory potential synthetic elements and genomic sequences are both impacted by the quality of the TFBS. The majority of synthetic elements, with consensus binding sites, drive expression above basal promoter levels, while genomic sequences with high scoring sites, ie: closer to the consensus sequence, are more likely to be active. The relationship between activity and TFBS scores is consistent with recent observations, specifically for the recruitment of ESRRB to areas enriched for OCT4, SOX2, and NANOG motifs (Adachi et al. 2018).

As for the grammar of combinations of TFBSs for these four pluripotency factors, in the controlled context of synthetic elements, we found clear evidence for a dependency on the position and number of the binding sites. While each site has a strong additive contribution to expression, consistent with the billboard model, we also uncovered evidence for some position-specific effects, best aligned with the enhanceosome model. The spacing and affinities of sites in synthetic elements were fixed, and the constant spacer sequences between sites reduced the likelihood of the presence of additional high quality TFBSs for other factors, constraints that are not representative of combinations of these sites in the genome. Indeed, when variable predicted affinity and distances between TFBSs is allowed, even with similar patterns of binding sites and evidence for activity dependent on one or more of those sites, the grammar learned from synthetic elements appears to have less of an impact on driving relative activity for genomic sequences. We found that many different arrangements of pluripotency sites can result in a high activity for genomic sequences, a prediction of the billboard model. However, because most genomic sequences were inactive, additional features, excluding identity and position, are also required to produce a sequence that can drive high activity. Surprisingly, active genomic sequences do not appear to have TFBSs for one or few additional factors that might point to a simple billboard model explanation for the patterns of expression observed. Instead, our modeling suggests that outside of the quality of the binding sites present, favorable distances between TFBSs and the ability to recruit TFs in their genomic context might contribute to the levels of expression observed for genomic sequences.

The literature demonstrates the utility of using synthetic elements to probe cis-regulatory grammar, understand interactions between factors, and predict true targets in the genome (Gertz, Siggia, and Cohen 2009; Cox, Surette, and Elowitz 2007). This work highlights the need for designing more nuanced synthetic elements, even when examining well-studied mammalian TFs, if we hope to predict the expected activity of patterns of TFBSs we observe in the genome. Additionally, we speculate that repression by some members of the pluripotency network might account for some of the seeming disparate behavior observed for similar combinations of OCT4, SOX2, KLF4 and ESRRB TFBSs from the genome. In particular, OCT4 has evidence of interactions with Polycomb and NuRD complex subunits in addition to physical interactions with Nanog (Jianlong Wang et al. 2006; Liang et al. 2008). The recruitment of repressive complexes and subsequent active repression would be difficult to distinguish from inactivity in current MPRA approaches. In the future, synthetic libraries of equally complex combinations of TFBS but where the predicted affinity and spacing are varied may better capture trends that allow us to understand the quantitative impact of these features in the genome and innovations in MPRA design to specifically detect active repression may allow for a more comprehensive understanding of the cis-regulatory landscape.

## Methods

### Library design

To generate a library that contained both synthetic and genomic elements we ordered a custom pool of 13,000 unique 150 bp oligonucleotides (oligos) from Agilent Technologies (Santa Clara, CA) through a limited licensing agreement. Each oligo in the SYN pool was 150 bp in length with the following sequence:

CTTCTACTACTAGGGCCCA[SEQ]AAGCTT[FILL]GAATTCTCTAGAC[BC]TGAGCTCTACATG CTAGTTCATG

where [SEQ] is a 40-80 bp synthetic element comprised of concatenated 20 bp building blocks of pluripotency sites, as described previously, with the fifth position of the KLF4 site changed to ‘T’ to facilitate cloning (Fiore and Cohen 2016). [FILL] is a random filler sequence of variable length to bring the total length of each sequence to 150 bp, and [BC] is a random 9 bp barcode. Synthetic elements were generated using a custom python script, generating all possible combinations of the pluripotency binding sites in both orientations, with no more than one of each site per sequence in lengths of two, three, and four building blocks (Supplemental Table 1). The sequence of each of the element is listed in Supplemental File 1. In total the SYN library has 624 unique synthetic elements. Each synthetic element is present in the pool eight times, each time with a different unique BC. There are also 112 oligos in the pool for cloning the basal promoter without any upstream element, each with a unique BC.

Genomic sequences were represented in the pool by 150 bp oligos with the following sequences:

GACTTACATTAGGGCCCGT[SEQ]AAGCTT[FILL]GAATTCTCTAGAC[BC]TGAGCTCGGACTA CGATACTG

Where [SEQ] is either a reference (gWT) or mutated (gMUT) genomic sequence of 81-82 bps. Reference gWT sequences were selected by choosing regions of the genome within 100 bps of previously identified ChIP-seq peaks for these four pluripotency factors (Chen et al. 2008). After excluding poorly sequenced and repetitive regions (The ENCODE Project Consortium 2012; Mouse Genome Sequencing Consortium et al. 2002), we scanned the remaining regions using FIMO with the four PWMs used previously to design the synthetic building blocks, with a p-value threshold of 1×10^−3^ (Grant, Bailey, and Noble 2011; Bailey et al. 2009; Fiore and Cohen 2016) Regions that contained more than one overlapping site identified by FIMO were excluded. Binding sites that were located less than 20bp from each other were then merged into a single genomic element using Bedtools (Quinlan and Hall 2010). Elements with no more than one of each site per element were then selected and expanded to 81-82 bp centered on the motifs. Expanded sequences were rescanned to confirm the presence of only three binding sites with the same threshold as used to originally scan the sequences. Sequences that contained restriction sites for were then removed from the library, leaving 407 genomic sequences with combinations of the OCT4, SOX2, KLF4, and/or ESSRB TFBSs (Supplemental Table 2).

We generated matched mutated sequences (gMUT) for each of the 407 gWT sequences by changing two positions in each motif from the highest information content base to the lowest information base for that position (Supplemental Figure S8). The reverse complement position and substitution was made for the reverse orientation of each motif. The mutated sequences were rescanned with all four original PWMs to confirm that no detectable pluripotency TFBSs remained, using FIMO with the same p-value threshold (1 × 10^−3^) as above.

In total the pool of oligos representing genomic sequences contained 407 wild type sequences (gWT) and the corresponding 407 gMUT sequences. The sequence of each of the element is listed in Supplemental File 2. Each of these 814 sequences were associated with eight unique BCs. The primers for gWT and gMUT sequences were identical so all subsequent steps for this library was performed in a single pool. There are also 112 oligos in the pool for cloning the basal promoter without any upstream element, each with a unique BC (Supplemental Table 3). The rest of the array contained sequences not used in this study.

### Cloning of plasmid libraries

The synthesized oligos were prepared as previously described (Kwasnieski et al. 2012; Fiore and Cohen 2016), except using primers Synthetic_FW-1 and Synthetic_Rev-2 with an annealing temperature of 55°C for the SYN library and primers Genomic_FW-1 and Genomic_Rev-1 with an annealing temperature of 53°C for the gWT/gMUT libraries (Supplemental Table 6). PCR products were purified from a polyacrylamide gel as described previously (White et al. 2013). Each library was cloned as described previously (Fiore and Cohen 2016), with a SYN element (SYN library) or either a gWT or gMUT sequence (gWT/gMUT library) cloned into the ApaI and SacI sites of plasmid pCF10. The pou5f1 basal promoter and dsRed reporter gene were amplified from pCF10 using primers CF121 and CF122, and inserted into the plasmid library pools from the previous step at the XbaI and HindIII sites. Digestion of the libraries with SpeI and subsequent size selection was omitted as the SYN library had less than 2% background and the combined gWT/gMUT library had less than 1% background in the final cloning step.

### Cell culture and transfection

RW4 mESCs were cultured as described previously (Xian, Werth, and Gottlieb 2005; C. T. L. Chen, Gottlieb, and Cohen 2008) on 2% gelatin coated plates in standard media (DMEM, 10% fetal bovine serum, 10% newborn calf serum, nucleoside supplement, 1000 U/ml leukemia inhibitory factor (LIF), and 0.1 uM B-mercaptoethanol). Approximately 1 millions cells at 100% estimated viability were seeded into 6-well plates 24 hours prior to transfection. The SYN library and combined gWT/gMUT were transfected in parallel using 10 uL Lipofectamine 2000 (Life Technologies, Carlsbad, CA), 3 ug of plasmid library, and 0.3 ug CF128 (a GFP control plasmid) per well, as described previously (Fiore and Cohen 2016). Four biological replicates of each library pool, the SYN plasmid pool or combined gWT/gMUT plasmid pool, were transfected and the plates were passaged 6 hours post-transfection. For three replicates of each library pool, RNA was extracted 24 hours post-transfection from approximately 9 million cells per replicate, using the PureLink RNA mini kit (Life Technologies, Carlsbad, CA) with the fourth transfection replicate reserved for estimating transfection efficiency via fluorescent microscopy and staining for alkaline phosphatase (AP) activity, a universal pluripotency marker (Singh et al. 2012).

### Massively Parallel Reporter Gene Assay

Massively parallel reporter gene assays were used to measure the activity of each CRE as described previously (Fiore and Cohen 2016; Mogno, Kwasnieski, and Cohen 2013). Briefly, we used Illumina NextSeq(tm) (San Deigo, CA) sequencing of both the RNA and original plasmid DNA pool, removing excess DNA from the RNA pool using TURBO DNA-free kit (Life Technologies, Carlsbad, CA). cDNA was then prepared using SuperScript RT III (Life Technologies, Carlsbad, CA) with oligo dT primers. Both the cDNA and the plasmid DNA pool were amplified using primers CF150 and CF151b (Supplemental Table 6), for 21 cycles. The PCR amplification products were digested using XbaI and XhoI (New England Biolabs, Ipswich, MA), ligating the resulting digestion products to custom Illumina adapter sequences, P1_XbaI_X (where X is 1 through 8, with in-line multiplexing BC sequences) to the 5’ overhang and PE2_SIC69_SalI on the 3’ XhoI overhang, each of which is comprised of annealed forward (F) and reverse (R) strands (Supplemental Table 6). An enrichment PCR with primers CF52 and CF53 was then used (Supplemental Table 6), and the resulting products were mixed at equal concentration and sequenced on one NextSeq lane.

Sequencing reads were filtered to ensure that the BC sequence perfectly matched the expected sequence. For the SYN library, this resulted in 40 million reads combined for the three demultiplexed RNA samples (P1_XbaI_1, P1_XbaI_2, P1_XbaI_3; 12.7-13.5 million each), and 19.7 million reads for the DNA library sample (P1_XbaI_7). For the combined gWT/gMUT libraries, this resulted in approximately 37 million reads combined for the three demultiplexed RNA samples (P1_XbaI_4, P1_XbaI_5, P1_XbaI_6; 9.4-16 million each), and 19.6 million reads for the DNA library sample (P1_XbaI_8). For each library, BCs that had less than 3 raw counts in any RNA replicate or less than 10 raw counts in the DNA sample were removed before proceeding with downstream analyses.

Expression normalization was performed by first calculating reads per million (RPM) per BC for each replicate for both the SYN library (R^2^ = 0.982-0.987; Figure S1A) and the combined gWT/gMUT library (R^2^ = 0.96-0.98; Figure S1B). For each BC, expression was calculated by dividing the RPMs in each RNA replicate by the DNA pool RPMs for that BC. Normalizing by DNA RPMs successfully removed the impact of the representation of the construct in the original pool as the calculated expression has no correlation with the DNA counts for both the SYN library (R^2^ = 0.02-0.02; Figure S1C) and the combined gWT/gMUT (R^2^ = 0.0005-0.008; Figure S1D). Within each biological replicate the BCs corresponding to each synthetic element (SYN) or genomic sequence (gWT/gMUT) were averaged and then normalized by basal mean expression in that replicate. These normalized expression values were then averaged across biological replicates. All downstream analyses were performed in R version 3.3.3 and plotted with ggplot2 version 2.2.1. Expression summaries per replicate are reported in Supplemental File 3 for the SYN library and Supplemental File 4 for the gWT/gMUT library.

### Predicted Occupancy

Custom code, based on Zhao & Stormo’s BEEML algorithm, was used to compare sequences of interest to a provided Energy Weight Matrix (EWM) at a set protein concentration (mu) and output a predicted occupancy for that TF (Zhao and Stormo 2011) as in White et al. 2013. Briefly, an energy landscape (EWM score) is calculated by comparing all n-mers of each sequence, where n = length of provided motif, to the matrix to generate an array of individual base scores for the forward and reverse orientation of the sequence. Occupancy is then predicted using equation 3 for binding probability at equilibrium, (1/ (1 + e^(ΔG – μ)^)). Position Frequency Matrices equivalent to the PWMs used for both SYN building block design and for scanning the mouse genome were used to generate EWMs, using the formula RT * ln(F req(Base ^^^ consensus)/F req(Base ^^^ i)) to convert the frequency of each base at each position i to a pseudo **Δ Δ**G values for each factor (White et al. 2013). Predicted occupancy (P(Occ)) for the 3-mer SYN elements was calculated for different assumed protein concentrations (mu = 0.5, 1, 2, 4, 5, 8, 10, 12) to determine at what point the SYN elements are predicted to be saturated, where P(Occ) ≅ 3 for each SYN element, i.e.: approaching 1 for each TFBS in the sequence. SYN elements were saturated by each of the four pluripotency factors at mu=8 with the exception of the shorter Oct4 motif, which reached saturation at mu=10. Occupancy of gWT and gMUT sequences was predicted for gWT and gMUT at an assumed high protein concentration of mu=8 for Sox2, Klf4, Esrrb, and mu =10 for Oct4, consistent with the role of these factors in mESCs. The predicted occupancy of each factor for matched gMUT sequences are reported in Supplemental File 8 as a feature of gWT sequences.

### iRF models

We built iterative Random Forest (iRF) models to classify our data using the R package iRF (version 2.0.0) from Basu et al. 2018. To run the software a model is initialized with 1/p weights for each of p features to be included in fitting the model. In each iteration, p features are reweighted by their Gini Importance (w^k^), a measure that is calculated by how purely a node, split by feature, separates the classes (Menze et al. 2009; Louppe et al. 2013). Default settings were used for model training, with four iterations of reweighting p features specified for each model as indicated in Supplemental Tables 5 & 6.

Synthetic data was split into training and test sets by randomly subsetting 50% of the total SYN elements (total n=407). Mean normalized expression was the response variable for model fitting for the synthetic models (see Supplemental File 7 for feature annotations for SYN elements). Four iterations of model fitting on training data was used.

Genomic data was split into training and test sets by randomly subsetting 50% of the total gWT/gMUT intervals (total n= 624). Classification as ‘active’, 1 if mean normalized gWT expression was greater than or equal to the 3rd quartile and ‘inactive’, 0 if mean normalized gWT expression was less than the 3rd quartile (cutoff value = 1.983), was the response variable for model fitting (see Supplemental File 8 for feature annotations and response values for gWT sequences). Four iterations of model fitting on training data was used.

### gkm-SVM

We used a gapped k-mer Support Vector Machine (gkm-SVM) to search for gapped k-mers that distinguish between highly active and inactive genomic sequences (Ghandi et al. 2016). We subset sequences from the gWT library into top 25% (high) and bottom 25% (low) based on expression data for a total of 101 positive and 101 negative intervals for the training set. FASTA sequences were then generated from the mm10 reference genome (Bioconductor, BioMart) for each region. We then used the gkm-SVM R package to classify high vs. low sequences (Ghandi et al. 2016). Word length (L) values of 6 (gap=2), 8 (gap=2), & 12 (gap=6), were tested with cross validation. Default settings were used for other function options.Three-fold cross validation was chosen due to the the amount of structure in the data, with combinations of OSK binding sites overrepresented in positive training sequences (See Figure S3). The best average performance on training data as evaluated by AUCs was the model trained with parameters of L= 8 and gap= 2. The final gkmer-SVM model includes approximately 1 million unique kmers (See Supplemental File 5 for full kmer list and weights).

### Other analysis and data sources

All genome coordinates from previous mouse genome builds were converted to mm10 using the UCSC liftover tool (Kuhn, Haussler, and Kent 2013). Binding matrices for SOX2, OCT4, KLF4, ESRRB were as previously reported (Fiore and Cohen 2016). The Bedtools suite (version 2.20) was used for manipulations and analysis of bed files (Quinlan and Hall 2010).

## Data Access

Raw sequencing data for SYN library and gWT/gMUT library can be found under SRA accession number SRP153192. Processed sequencing data reported in this paper, including expression per replicate and demultiplexed barcode counts, are included as Supplemental File 3 & Supplemental File 9 for the SYN library and Supplemental File 4 & Supplemental File 10 for the gWT/gMUT library.

## Acknowledgements

We thank members of the Cohen Lab for critical reading and feedback, particularly Michael White, Max Staller and Hemangi Chaudhari for helpful discussion over the course of the project, and Jessica Hoisington-Lopez from the DNA Sequencing Innovation Lab for assistance with high-throughput sequencing. This work is supported by a grant from the National Institutes of Health, R01 GM092910 to B.A.C.

## Author Contributions

D.M.K and B.A.C. designed the experiments. D.M.K and B.M. collected the data. D.M.K analyzed the data. D.M.K and B.A.C wrote the manuscript.

## References

Adachi, Kenjiro, Wolfgang Kopp, Guangming Wu, Sandra Heising, Boris Greber, Martin Stehling, Marcos J. Araúzo-Bravo, et al. 2018. “Esrrb Unlocks Silenced Enhancers for Reprogramming to Naive Pluripotency.” Cell Stem Cell, June. https://doi.org/10.1016/j.stem.2018.05.020.

Bailey, Timothy L., Mikael Boden, Fabian a. Buske, Martin Frith, Charles E. Grant, Luca Clementi, Jingyuan Ren, Wilfred W. Li, and William S. Noble. 2009. “MEME SUITE: Tools for Motif Discovery and Searching.” Nucleic Acids Research 37 (Web Server issue): W202–8. https://doi.org/10.1093/nar/gkp335.

Basu, Sumanta, Karl Kumbier, James B. Brown, and Bin Yu. 2018. “Iterative Random Forests to Discover Predictive and Stable High-Order Interactions.” Proceedings of the National Academy of Sciences of the United States of America 115 (8): 1943–48. https://doi.org/10.1073/pnas.1711236115.

Chambers, Ian, and Simon R. Tomlinson. 2009. “The Transcriptional Foundation of Pluripotency.” Development 136: 2311–22. https://doi.org/10.1242/dev.024398.

Chaudhari, Hemangi G., and Barak A. Cohen. 2018. “Local Sequence Features That Influence AP-1 Cis-Regulatory Activity.” Genome Research 28 (2): 171–81. https://doi.org/10.1101/gr.226530.117.

Chen, Chih-Yu, Quaid Morris, and Jennifer a. Mitchell. 2012. “Enhancer Identification in Mouse Embryonic Stem Cells Using Integrative Modeling of Chromatin and Genomic Features.” BMC Genomics 13 (1): 152. https://doi.org/10.1186/1471-2164-13-152.

Chen, Christina T. L., David I. Gottlieb, and Barak A. Cohen. 2008. “Ultraconserved Elements in the Olig2 Promoter.” PloS One 3 (12): e3946. https://doi.org/10.1371/journal.pone.0003946.

Chen, Xi, and Hemant Ishwaran. 2012. “Random Forests for Genomic Data Analysis.” Genomics 99 (6): 323–29. https://doi.org/10.1016/j.ygeno.2012.04.003.

Chen, Xi, Han Xu, Ping Yuan, Fang Fang, Mikael Huss, Vinsensius B. Vega, Eleanor Wong, et al. 2008. “Integration of External Signaling Pathways with the Core Transcriptional Network in Embryonic Stem Cells.” Cell 133: 1106–17. https://doi.org/10.1016/j.cell.2008.04.043.

Chen, X., V. B. Vega, and H-H Ng. 2008. “Transcriptional Regulatory Networks in Embryonic Stem Cells.” Cold Spring Harbor Symposia on Quantitative Biology 73: 203–9. https://doi.org/10.1101/sqb.2008.73.026.

Cox, Robert Sidney, 3rd, Michael G. Surette, and Michael B. Elowitz. 2007. “Programming Gene Expression with Combinatorial Promoters.” Molecular Systems Biology 3 (November): 145. https://doi.org/10.1038/msb4100187.

Dunn, S-J, G. Martello, B. Yordanov, S. Emmott, and a. G. Smith. 2014. “Defining an Essential Transcription Factor Program for Naïve Pluripotency.” Science 344: 1156–60. https://doi.org/10.1126/science.1248882.

Evans, Nicole C., Christina I. Swanson, and Scott Barolo. 2012. Sparkling Insights into Enhancer Structure, Function, and Evolution. 1st ed. Vol. 98. Elsevier Inc. https://doi.org/10.1016/B978-0-12-386499-4.00004-5.

Feng, Bo, Jianming Jiang, Petra Kraus, Jia-Hui Ng, Jian-Chien Dominic Heng, Yun-Shen Chan, Lai-Ping Yaw, et al. 2009. “Reprogramming of Fibroblasts into Induced Pluripotent Stem Cells with Orphan Nuclear Receptor Esrrb.” Nature Cell Biology 11 (2): 197–203. https://doi.org/10.1038/ncb1827.

Fiore, Chris, and Barak A. Cohen. 2016. “Interactions between Pluripotency Factors Specify Cis-Regulation in Embryonic Stem Cells.” Genome Research 26 (6): 778–86. https://doi.org/10.1101/gr.200733.115.

Fisher, William W., Jingyi Jessica Li, Ann S. Hammonds, James B. Brown, Barret D. Pfeiffer, Richard Weiszmann, Stewart MacArthur, et al. 2012. “DNA Regions Bound at Low Occupancy by Transcription Factors Do Not Drive Patterned Reporter Gene Expression in Drosophila.” Proceedings of the National Academy of Sciences of the United States of America 109 (52): 21330–35. https://doi.org/10.1073/pnas.1209589110.

Fletez-Brant, Christopher, Dongwon Lee, Andrew S. McCallion, and Michael A. Beer. 2013. “Kmer-SVM: A Web Server for Identifying Predictive Regulatory Sequence Features in Genomic Data Sets.” Nucleic Acids Research 41 (Web Server issue): W544–56. https://doi.org/10.1093/nar/gkt519.

Gabut, Mathieu, Payman Samavarchi-Tehrani, Xinchen Wang, Valentina Slobodeniuc, Dave O’Hanlon, Hoon Ki Sung, Manuel Alvarez, et al. 2011. “An Alternative Splicing Switch Regulates Embryonic Stem Cell Pluripotency and Reprogramming.” Cell 147 (1): 132–46. https://doi.org/10.1016/j.cell.2011.08.023.

Gertz, Jason, Eric D. Siggia, and Barak A. Cohen. 2009. “Analysis of Combinatorial Cis-Regulation in Synthetic and Genomic Promoters.” Nature 457 (7226): 215–18. https://doi.org/10.1038/nature07521.

Ghandi, Mahmoud, Dongwon Lee, Morteza Mohammad-Noori, and Michael A. Beer. 2014. “Enhanced Regulatory Sequence Prediction Using Gapped K-Mer Features.” PLoS Computational Biology 10 (7): e1003711. https://doi.org/10.1371/journal.pcbi.1003711.

Ghandi, Mahmoud, Morteza Mohammad-Noori, Narges Ghareghani, Dongwon Lee, Levi Garraway, and Michael A. Beer. 2016. “gkmSVM: An R Package for Gapped-Kmer SVM.” Bioinformatics 32 (14): 2205–7. https://doi.org/10.1093/bioinformatics/btw203.

Giorgetti, Luca, Trevor Siggers, Guido Tiana, Greta Caprara, Samuele Notarbartolo, Teresa Corona, Manolis Pasparakis, Paolo Milani, Martha L. Bulyk, and Gioacchino Natoli. 2010. “Noncooperative Interactions between Transcription Factors and Clustered DNA Binding Sites Enable Graded Transcriptional Responses to Environmental Inputs.” Molecular Cell 37 (3): 418–28. https://doi.org/10.1016/j.molcel.2010.01.016.

Grant, Charles E., Timothy L. Bailey, and William Stafford Noble. 2011. “FIMO: Scanning for Occurrences of a given Motif.” Bioinformatics 27 (7): 1017–18. https://doi.org/10.1093/bioinformatics/btr064.

Grossman, Sharon R., Xiaolan Zhang, Li Wang, Jesse Engreitz, Alexandre Melnikov, Peter Rogov, Ryan Tewhey, et al. 2017. “Systematic Dissection of Genomic Features Determining Transcription Factor Binding and Enhancer Function.” Proceedings of the National Academy of Sciences of the United States of America 114 (7): E1291–1300. https://doi.org/10.1073/pnas.1621150114.

Hare, Emily E., Brant K. Peterson, and Michael B. Eisen. 2008. “A Careful Look at Binding Site Reorganization in the Even-Skipped Enhancers of Drosophila and Sepsids.” PLoS Genetics 4 (11): 1–5. https://doi.org/10.1371/journal.pgen.1000268.

Hare, Emily E., Brant K. Peterson, Venky N. Iyer, Rudolf Meier, and Michael B. Eisen. 2008. “Sepsid Even-Skipped Enhancers Are Functionally Conserved in Drosophila despite Lack of Sequence Conservation.” PLoS Genetics 4 (6): e1000106. https://doi.org/10.1371/journal.pgen.1000106.

Huang, Jinyan, Taotao Chen, Xiaosong Liu, Jing Jiang, Jinsong Li, Dangsheng Li, X. Shirley Liu, Wei Li, Jiuhong Kang, and Gang Pei. 2009. “More Synergetic Cooperation of Yamanaka Factors in Induced Pluripotent Stem Cells than in Embryonic Stem Cells.” Cell Research 19 (10): 1127–38. https://doi.org/10.1038/cr.2009.106.

Jauch, Ralf, Calista Keow Leng Ng, Kumar Singh Saikatendu, Raymond C. Stevens, and Prasanna R. Kolatkar. 2008. “Crystal Structure and DNA Binding of the Homeodomain of the Stem Cell Transcription Factor Nanog.” Journal of Molecular Biology 376 (3): 758–70. https://doi.org/10.1016/j.jmb.2007.11.091.

Junion, Guillaume, Mikhail Spivakov, Charles Girardot, Martina Braun, E. Hilary Gustafson, Ewan Birney, and Eileen E. M. Furlong. 2012. “A Transcription Factor Collective Defines Cardiac Cell Fate and Reflects Lineage History.” Cell 148 (3): 473–86. https://doi.org/10.1016/j.cell.2012.01.030.

Kheradpour, Pouya, Jason Ernst, Alexandre Melnikov, Peter Rogov, Li Wang, Xiaolan Zhang, Jessica Alston, Tarjei S. Mikkelsen, and Manolis Kellis. 2013. “Systematic Dissection of Regulatory Motifs in 2000 Predicted Human Enhancers Using a Massively Parallel Reporter Assay.” Genome Research 23 (5): 800–811. https://doi.org/10.1101/gr.144899.112.

Kuhn, Robert M., David Haussler, and W. James Kent. 2013. “The UCSC Genome Browser and Associated Tools.” Briefings in Bioinformatics 14 (2): 144–61. https://doi.org/10.1093/bib/bbs038.

Kulkarni, Meghana M., and David N. Arnosti. 2003. “Information Display by Transcriptional Enhancers.” Development 130: 6569–75. https://doi.org/10.1242/dev.00890.

Kwasnieski, Jamie C., Ilaria Mogno, Connie A. Myers, Joseph C. Corbo, and Barak A. Cohen. 2012. “Complex Effects of Nucleotide Variants in a Mammalian Cis -Regulatory Element.” Proceedings of the National Academy of Sciences of the United States of America 109 (47): 19498–503. https://doi.org/10.1073/pnas.1210678109/-/DCSupplemental.www.pnas.org/cgi/doi/10.1073/pnas.1210678109.

Liang, Jiancong, Ma Wan, Yi Zhang, Peili Gu, Huawei Xin, Sung Yun Jung, Jun Qin, et al. 2008. “Nanog and Oct4 Associate with Unique Transcriptional Repression Complexes in Embryonic Stem Cells.” Nature Cell Biology 10 (6): 731–39. https://doi.org/10.1038/ncb1736.

Liu, Xiaosong, Jinyan Huang, Taotao Chen, Ying Wang, Shunmei Xin, Jian Li, Gang Pei, and Jiuhong Kang. 2008. “Yamanaka Factors Critically Regulate the Developmental Signaling Network in Mouse Embryonic Stem Cells.” Cell Research 18: 1177–89. https://doi.org/10.1038/cr.2008.309.

Louppe, Gilles, Louis Wehenkel, Antonio Sutera, and Pierre Geurts. 2013. “Understanding Variable Importances in Forests of Randomized Trees.” In Advances in Neural Information Processing Systems 26, edited by C. J. C. Burges, L. Bottou, M. Welling, Z. Ghahramani, and K. Q. Weinberger, 431–39.

Curran Associates, Inc. http://papers.nips.cc/paper/4928-understanding-variable-importances-in-forests-of-randomized-trees.pdf.

Ludwig, M. Z., C. Bergman, N. H. Patel, and M. Kreitman. 2000. “Evidence for Stabilizing Selection in a Eukaryotic Enhancer Element.” Nature 403 (6769): 564–67. https://doi.org/10.1038/35000615.

Menze, Bjoern H., B. Michael Kelm, Ralf Masuch, Uwe Himmelreich, Peter Bachert, Wolfgang Petrich, and Fred A. Hamprecht. 2009. “A Comparison of Random Forest and Its Gini Importance with Standard Chemometric Methods for the Feature Selection and Classification of Spectral Data.” BMC Bioinformatics 10 (July): 213. https://doi.org/10.1186/1471-2105-10-213.

Mogno, Ilaria, Jamie C. Kwasnieski, and Barak A. Cohen. 2013. “Massively Parallel Synthetic Promoter Assays Reveal the in Vivo Effects of Binding Site Variants.” Genome Research 23 (11): 1908–15. https://doi.org/10.1101/gr.157891.113.

Mouse Genome Sequencing Consortium, Robert H. Waterston, Kerstin Lindblad-Toh, Ewan Birney, Jane Rogers, Josep F. Abril, Pankaj Agarwal, et al. 2002. “Initial Sequencing and Comparative Analysis of the Mouse Genome.” Nature 420 (6915): 520–62. https://doi.org/10.1038/nature01262.

Mulas, Carla, Gloryn Chia, Kenneth Alan Jones, Andrew Christopher Hodgson, Giuliano Giuseppe Stirparo, and Jennifer Nichols. 2018. “Oct4 Regulates the Embryonic Axis and Coordinates Exit from Pluripotency and Germ Layer Specification in the Mouse Embryo.” Development 145 (12). https://doi.org/10.1242/dev.159103.

Niwa, Hitoshi. 2014. “The Pluripotency Transcription Factor Network at Work in Reprogramming.” Current Opinion in Genetics & Development 28: 25–31. https://doi.org/10.1016/j.gde.2014.08.004.

Pan, Guangjin, and James a. Thomson. 2007. “Nanog and Transcriptional Networks in Embryonic Stem Cell Pluripotency.” Cell Research 17: 42–49. https://doi.org/10.1038/sj.cr.7310125.

Panne, Daniel. 2008. “The Enhanceosome.” Current Opinion in Structural Biology 18: 236–42. https://doi.org/10.1016/j.sbi.2007.12.002.

Patwardhan, Rupali P., Joseph B. Hiatt, Daniela M. Witten, Mee J. Kim, Robin P. Smith, Dalit May, Choli Lee, et al. 2012. “Massively Parallel Functional Dissection of Mammalian Enhancers in Vivo.” Nature Biotechnology 30 (3): 265–70. https://doi.org/10.1038/nbt.2136.

Quinlan, Aaron R., and Ira M. Hall. 2010. “BEDTools: A Flexible Suite of Utilities for Comparing Genomic Features.” Bioinformatics 26 (6): 841–42. https://doi.org/10.1093/bioinformatics/btq033.

Reményi, Attila, Katharina Lins, L. Johan Nissen, Rolland Reinbold, Hans R. Schöler, and Matthias Wilmanns. 2003. “Crystal Structure of a POU/HMG/DNA Ternary Complex Suggests Differential Assembly of Oct4 and Sox2 on Two Enhancers.” Genes & Development 17 (16): 2048–59. https://doi.org/10.1101/gad.269303.

Reményi, Attila, Hans R. Schöler, and Matthias Wilmanns. 2004. “Combinatorial Control of Gene Expression.” Nature Structural & Molecular Biology 11 (9): 812–15. https://doi.org/10.1038/nsmb820.

Sandelin, Albin, Wynand Alkema, Pär Engström, Wyeth W. Wasserman, and Boris Lenhard. 2004. “JASPAR: An Open-Access Database for Eukaryotic Transcription Factor Binding Profiles.” Nucleic Acids Research 32 (Database issue): D91–94. https://doi.org/10.1093/nar/gkh012.

Segal, Eran, Tali Raveh-Sadka, Mark Schroeder, Ulrich Unnerstall, and Ulrike Gaul. 2008. “Predicting Expression Patterns from Regulatory Sequence in Drosophila Segmentation.” Nature 451 (7178): 535–40. https://doi.org/10.1038/nature06496.

Singh, Upinder, Rene H. Quintanilla, Scott Grecian, Kyle R. Gee, Mahendra S. Rao, and Uma Lakshmipathy. 2012. “Novel Live Alkaline Phosphatase Substrate for Identification of Pluripotent Stem Cells.” Stem Cell Reviews 8 (3): 1021–29. https://doi.org/10.1007/s12015-012-9359-6.

Spitz, François, and Eileen E. M. Furlong. 2012. “Transcription Factors: From Enhancer Binding to Developmental Control.” Nature Reviews. Genetics 13 (9): 613–26. https://doi.org/10.1038/nrg3207.

Takahashi, Kazutoshi, and Shinya Yamanaka. 2006. “Induction of Pluripotent Stem Cells from Mouse Embryonic and Adult Fibroblast Cultures by Defined Factors.” Cell 126 (4): 663–76. https://doi.org/10.1016/j.cell.2006.07.024.

The ENCODE Project Consortium. 2012. “An Integrated Encyclopedia of DNA Elements in the Human Genome.” Nature 489 (September): 57. https://doi.org/10.1038/nature11247.

Uhl, Juli D., Arya Zandvakili, and Brian Gebelein. 2016. “A Hox Transcription Factor Collective Binds a Highly Conserved Distal-Less Cis-Regulatory Module to Generate Robust Transcriptional Outcomes.” PLoS Genetics 12 (4): e1005981. https://doi.org/10.1371/journal.pgen.1005981.

Ulirsch, Jacob C., Satish K. Nandakumar, Li Wang, Felix C. Giani, Xiaolan Zhang, Peter Rogov, Alexandre Melnikov, et al. 2016. “Systematic Functional Dissection of Common Genetic Variation Affecting Red Blood Cell Traits.” Cell 165 (6): 1530–45. https://doi.org/10.1016/j.cell.2016.04.048.

Visel, Axel, Matthew J. Blow, Zirong Li, Tao Zhang, Jennifer A. Akiyama, Amy Holt, Ingrid Plajzer-frick, et al. 2009. “ChIP-Seq Accurately Predicts Tissue-Specific Activity of Enhancers.” Nature 457 (7231): 854–58. https://doi.org/10.1038/nature07730.

Wang, Jianlong, Sridhar Rao, Jianlin Chu, Xiaohua Shen, Dana N. Levasseur, Thorold W. Theunissen, and Stuart H. Orkin. 2006. “A Protein Interaction Network for Pluripotency of Embryonic Stem Cells.” Nature 444 (November): 364–68. https://doi.org/10.1038/nature05284.

Wang, Jie, Jiali Zhuang, Sowmya Iyer, Xin-Ying Lin, Melissa C. Greven, Bong-Hyun Kim, Jill Moore, et al. 2013. “Factorbook.org: A Wiki-Based Database for Transcription Factor-Binding Data Generated by the ENCODE Consortium.” Nucleic Acids Research 41 (Database issue): D171–76. https://doi.org/10.1093/nar/gks1221.

Wang, Jie, Jiali Zhuang, Sowmya Iyer, Xinying Lin, Troy W. Whitfield, Melissa C. Greven, Brian G. Pierce, et al. 2012. “Sequence Features and Chromatin Structure around the Genomic Regions Bound by 119 Human Transcription Factors.” Genome Research 22 (9): 1798–1812. https://doi.org/10.1101/gr.139105.112.

White, Michael A. 2015. “Understanding How Cis-Regulatory Function Is Encoded in DNA Sequence Using Massively Parallel Reporter Assays and Designed Sequences.” Genomics 106 (3): 165–70. https://doi.org/10.1016/j.ygeno.2015.06.003.

White, Michael A., Jamie C. Kwasnieski, Connie A. Myers, Susan Q. Shen, Joseph C. Corbo, and Barak A. Cohen. 2016. “A Simple Grammar Defines Activating and Repressing Cis-Regulatory Elements in Photoreceptors.” Cell Reports 17 (5): 1247–54. https://doi.org/10.1016/j.celrep.2016.09.066.

White, Michael A., Connie A. Myers, Joseph C. Corbo, and Barak A. Cohen. 2013. “Massively Parallel in Vivo Enhancer Assay Reveals That Highly Local Features Determine the Cis -Regulatory Function of ChIP-Seq Peaks.” Proceedings of the National Academy of Sciences 110 (29): 11952–57. https://doi.org/10.1073/pnas.1307449110/-/DCSupplemental.www.pnas.org/cgi/doi/10.1073/pnas.1307449110.

Williams, David C., Mengli Cai, and G. Marius Clore. 2004. “Molecular Basis for Synergistic Transcriptional Activation by Oct1 and Sox2 Revealed from the Solution Structure of the 42-kDa Oct1 · Sox2 · Hoxb1-DNA Ternary Transcription Factor Complex.” The Journal of Biological Chemistry 279 (2): 1449–57. https://doi.org/10.1074/jbc.M309790200.

Xian, Hai-Qing, Kelly Werth, and David I. Gottlieb. 2005. “Promoter Analysis in ES Cell-Derived Neural Cells.” Biochemical and Biophysical Research Communications 327 (1): 155–62. https://doi.org/10.1016/j.bbrc.2004.11.149.

Yie, J., K. Senger, and D. Thanos. 1999. “Mechanism by Which the IFN-Beta Enhanceosome Activates Transcription.” Proceedings of the National Academy of Sciences of the United States of America 96 (Track II): 13108–13. https://doi.org/10.1073/pnas.96.23.13108.

Zhang, Xiaofei, Juan Zhang, Tao Wang, Miguel a. Esteban, and Duanqing Pei. 2008. “Esrrb Activates Oct4 Transcription and Sustains Self-Renewal and Pluripotency in Embryonic Stem Cells.” The Journal of Biological Chemistry 283: 35825–33. https://doi.org/10.1074/jbc.M803481200.

Zhao, Yue, and Gary D. Stormo. 2011. “Quantitative Analysis Demonstrates Most Transcription Factors Require Only Simple Models of Specificity.” Nature Biotechnology 29 (6): 480–83. https://doi.org/10.1038/nbt.1893.

